# A strain-specific inhibitor of receptor-bound HIV-1 targets a pocket near the fusion peptide and offers a template for drug design

**DOI:** 10.1101/2020.06.11.146654

**Authors:** Gabriel Ozorowski, Jonathan L. Torres, Diogo Santos-Martins, Stefano Forli, Andrew B. Ward

## Abstract

Disruption of viral fusion represents a viable, albeit under-explored, target for HIV therapeutics. While studying the receptor-bound envelope glycoprotein conformation by cryo-EM, we identified a pocket near the base of the trimer containing a bound detergent molecule and performed *in silico* drug screening using a library of drug-like and commercially available molecules. After down-selection, we solved cryo-EM structures that validated binding of two small molecule hits in very similar manners to the predicted binding poses, including interactions with aromatic residues within the fusion peptide. One of the molecules demonstrated low micromolar inhibition of the autologous virus by utilizing a very rare phenylalanine in the fusion peptide and stabilizing the surrounding region. This work demonstrates that small molecules can target the fusion process, providing a new target for anti-HIV therapeutics, and highlights the need to explore how fusion peptide sequence variations affect receptor-mediated conformational states across diverse HIV strains.

## Introduction

Despite advances in the characterization of HIV and treatment of infected individuals, both a functional cure and prophylactic vaccine are lacking (*1, 2*). This situation almost ensures that the number of global HIV infected individuals will continue to rise, even under the most aggressive efforts from the medical community that have partially succeeded in slowing down the annual rate of infection (https://www.who.int/hiv/data/en/). Viremia in HIV positive individuals can be well controlled using antiretroviral therapy (ART), which provides a relatively high quality of life by halting the progression to AIDS (*3*). Furthermore, proper ART decreases HIV transmission and will continue to have a major role in fighting the HIV pandemic. Current ART methods utilize small molecule drugs, however recently a new class of potential HIV therapeutics, broadly neutralizing antibodies, has also been shown to suppress viremia in infected individuals(*4*). The elicitation of such antibodies is the ultimate goal of HIV vaccine efforts, and the utility of recombinantly expressed versions of these antibodies for prophylaxis and ART continues to be heavily investigated. Approved ART drugs target either HIV-specific enzymes (reverse transcriptase, protease, integrase), HIV fusion, or HIV receptors/co-receptors (CD4, CCR5) (https://aidsinfo.nih.gov/understanding-hiv-aids/fact-sheets/21/58/fda-approved-hiv-medicines). Of the currently over 3 dozen FDA-approved HIV medicines, only 1, enfuvirtide, is a fusion inhibitor. Because viral fusion to the host cell is a necessary and conserved first step of HIV infection, the discovery of new inhibitors may lead to better antiretroviral therapies that are less prone to drug resistance.

HIV fusion is facilitated by the viral envelope glycoprotein (Env), a trimer of non-covalently linked heterodimers (gp120 and gp41)(*5, 6*). Binding of the receptor CD4 to gp120 triggers a series of conformational changes, including opening of the trimer, exposure of the co-receptor binding sites, and rearrangements of the gp41 helices(*5, 7, 8*). The N-terminal region of gp41 forms the fusion peptide (FP), which becomes sequestered during the initial steps of receptor binding by moving towards the trimer interior(*7*). After receptor and co-receptor binding the trimer is thought to undergo even more major conformational changes, such as gp120 shedding and the formation of the 6-helix bundle, eventually leading to the insertion of the FP into the host membrane and fusion with the viral membrane(*5*). The FDA-approved fusion inhibitor enfuvirtide is a peptide drug mimetic that resembles a portion of the HR2 helix of gp41 and is thought to disrupt one of the penultimate gp41 changes prior to membrane fusion(*9*). Other reported small molecules that target HIV fusion and have demonstrated inhibitory activity bind instead to the closed, pre-receptor engagement state of the Env trimer, such as candidate drug molecules that are based on the Bristol Myers Squibb inhibitor BMS-626529(*10*). In recent years, these molecules have generated excitement, with many published structures, safety and efficacy reports, as well as ongoing phase III clinical trials(*10, 11*). These molecules work by binding the pre-fusion, receptor-free states of Env and halt conformational changes associated with receptor binding(*12*).

Advances in cryo-EM, a plethora of anti-HIV antibodies, and synthetic ligands now provide tools for structural elucidation of various Env conformations. These transient states represent new targets for small molecules inhibitors, similar to what has been done for GPCRs(*13, 14*). Here, we identified a pocket in gp41 of our previous cryo-EM reconstruction of an early pre-fusion intermediate Env (CD4- and co-receptor mimic antibody-bound)(*7*) that is proximal to the FP and contains a bound detergent molecule used during cryo-EM grid preparation. Guided by this reference molecule and its interactions with residues lining the pocket, we performed *in silico* drug screening using a library of drug-like and commercially available small molecules. Through a combination of biophysical methods, including cryo-EM, we confirmed that two of the molecules specifically bound the pocket, in very similar manners to their predicted binding poses. One molecule in particular inhibited viral entry at low micromolar levels.

## Results

### A potentially druggable pocket forms near the FP after receptor binding

We previously reported cryo-EM maps of SOSIP in complex with b12 or CD4/17b that demonstrated a distinct and stable conformation of the fusion peptide and fusion peptide proximal region (FPPR) upon receptor- or antibody-induced trimer opening(*7*). In the original (C3-symmetric) CD4-bound structure we omitted the first 3 residues of the FP (A512-G514) from the atomic model due to local disorder, resulting in unassigned density within the vicinity of FP/FFPR. As an attempt to better resolve this region, the data was reprocessed using newer software (Relion 3.0(*15*), CryoSPARC version 2(*16*)), including template-based particle picking to extract more particles that may have been missed by the previous difference of Gaussians approach(*17*). Reprocessing resulted in well-resolved C3-symmetric and asymmetric reconstructions that each exceeded the FSC resolution estimate of the original map (C1: 3.6 Å, C3: 3.3 Å; EMD-8713 C3 map: 3.7 Å) and allowed for better interpretation of the FP/FPPR region (**Fig. S1, A to D, and table S1**). We attribute most of the improvement to an increase in the number of particles in the final reconstruction (nearly 4x that of the originally published map), which was streamlined by the template-based particle picker of CryoSPARC version 2(*16*).

The FP can now be fully modeled into the new C3-symmetry map and is almost completely resolved in 2 out of 3 protomers of the asymmetric map (**Fig. 1A**). Intriguingly, both of the new maps contain additional resolved density for a long and narrow small molecule proximal to the FP in all protomers, although in the asymmetric reconstruction this is less prominent in the protomer with a less-resolved FP (**Fig. 1B, and fig. S1E**). Because cryo-EM freezing techniques often include sub-critical micellar concentration (CMC) amounts of detergent to increase the number and tumbling of protein particles trapped over holes in vitreous ice, we hypothesized that the unassigned density could be the DDM (n-dodecyl-β-D-maltoside) used in our experiment. In fact, 2 full DDM molecules and 1 partial DDM molecule could be built and refined into the C1 map, while C3-symmetry averaging enhances the signal for DDM in all binding sites (**Fig. 1B, and fig. S1E**). The partial density for one of the three DDM molecules in the asymmetric reconstruction may be a result of sub-stoichiometric binding due to the low detergent concentration in the solution (approximately 2-3X molar excess of DDM to trimer), or intrinsic local asymmetry does not favor uniform binding. Recent cryo-EM reconstructions of Env SOSIP from the BG505 genotype in complex with sCD4 and a different co-receptor mimic antibody (E51) suggest asymmetry amongst the 3 protomers may be a naturally-occurring feature of Env, at least in the context of receptor-binding to the soluble, stabilized SOSIP construct (*18*). Another notable feature of the DDM-proximal residues (excluding the FP) is high conservation across HIV-genotypes (**Fig. 1, C and D**). Hence, we further explored this pocket as a potential site for small molecule inhibitors.

**Fig. 1.**
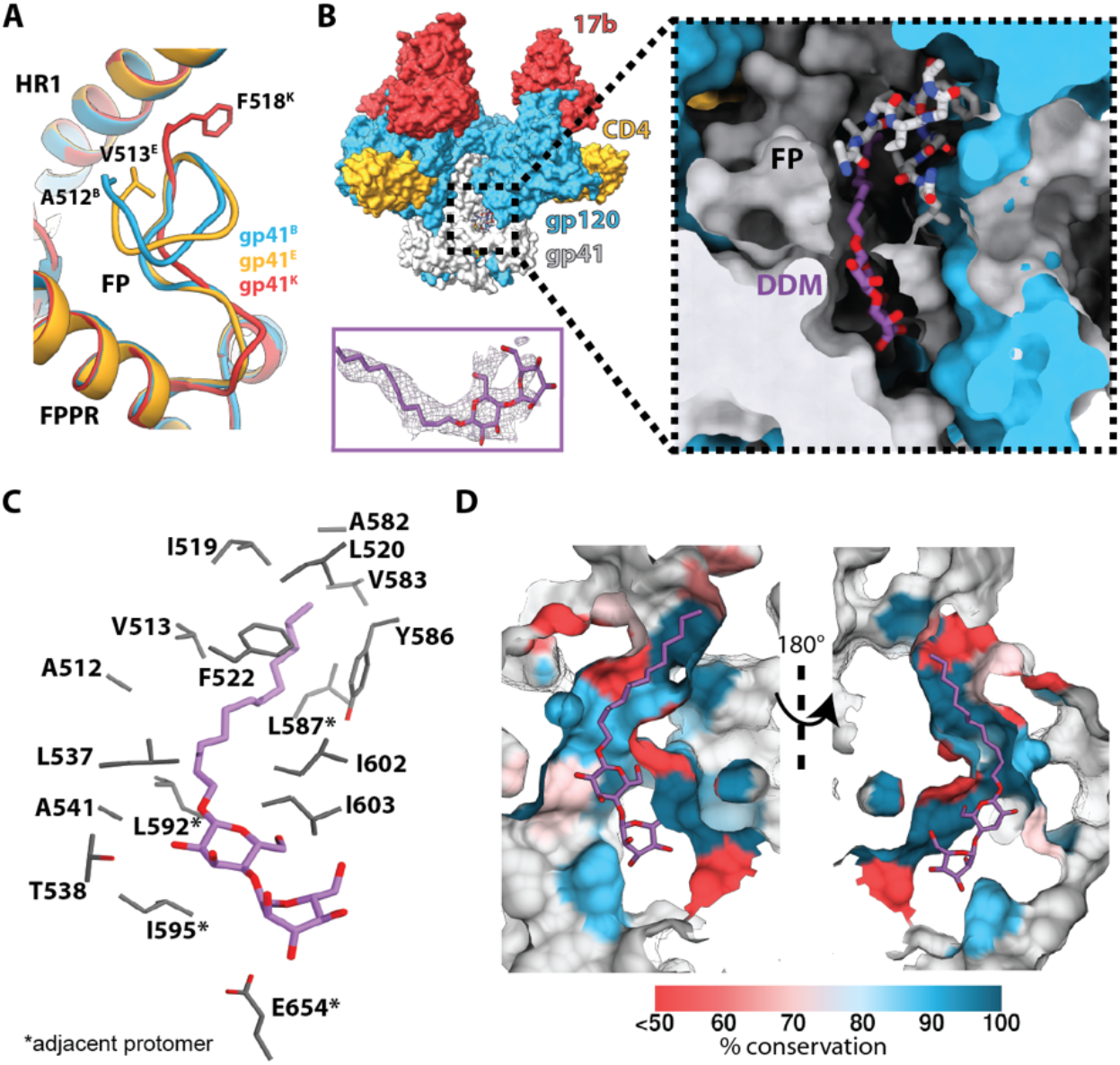
A detergent molecule binds a receptor-induced pocket in HIV-1 Env. (**A**) The fusion peptide adopts different conformations in the asymmetric reconstruction of CD4- and 17b-bound B41 SOSIP. Modeled N-terminal residues of each chain are labeled. (**B**) Relative location of the binding pocket (*left*), and greater details of the locations of the DDM-containing and FP-containing pockets (*right*). The density for DDM in the C3-symmetry map is shown as a side panel for reference. (**C**) Contact residues and (**D**) percent conservation of residues lining the DDM pocket.

### Using cryo-EM models for virtual screening

The refined coordinates of DDM were used to define a ligand binding pocket to conduct *in silico* virtual screening and identify other molecules that could potentially bind. To facilitate the process, we started with the higher resolution C3-symmetric model. AutoSite(*19*) software was used to analyze the protein structure and identify the location and the size of the optimal ligand volume at the DDM binding site (**Fig. 2A**). The docking box was centered on the AutoSite volume in one of the pockets (at the interface between gp120 chain A and gp41 chains B and M), then expanded to include the larger opening engaged by the maltose moieties (with orthogonal corners roughly located between A582 and Q658) (**Fig. 2A**). The resulting docking box was significantly larger than the reference ligand and the predicted optimal volume. This large box enabled exploration of extra hydrophilic interactions near the distal glucose ring, as well as to accommodate potential uncertainties associated with the coordinates in the cryo-EM model (e.g. decarboxylation of acidic side chains due to radiation damage(*20*)).

**Fig. 2.**
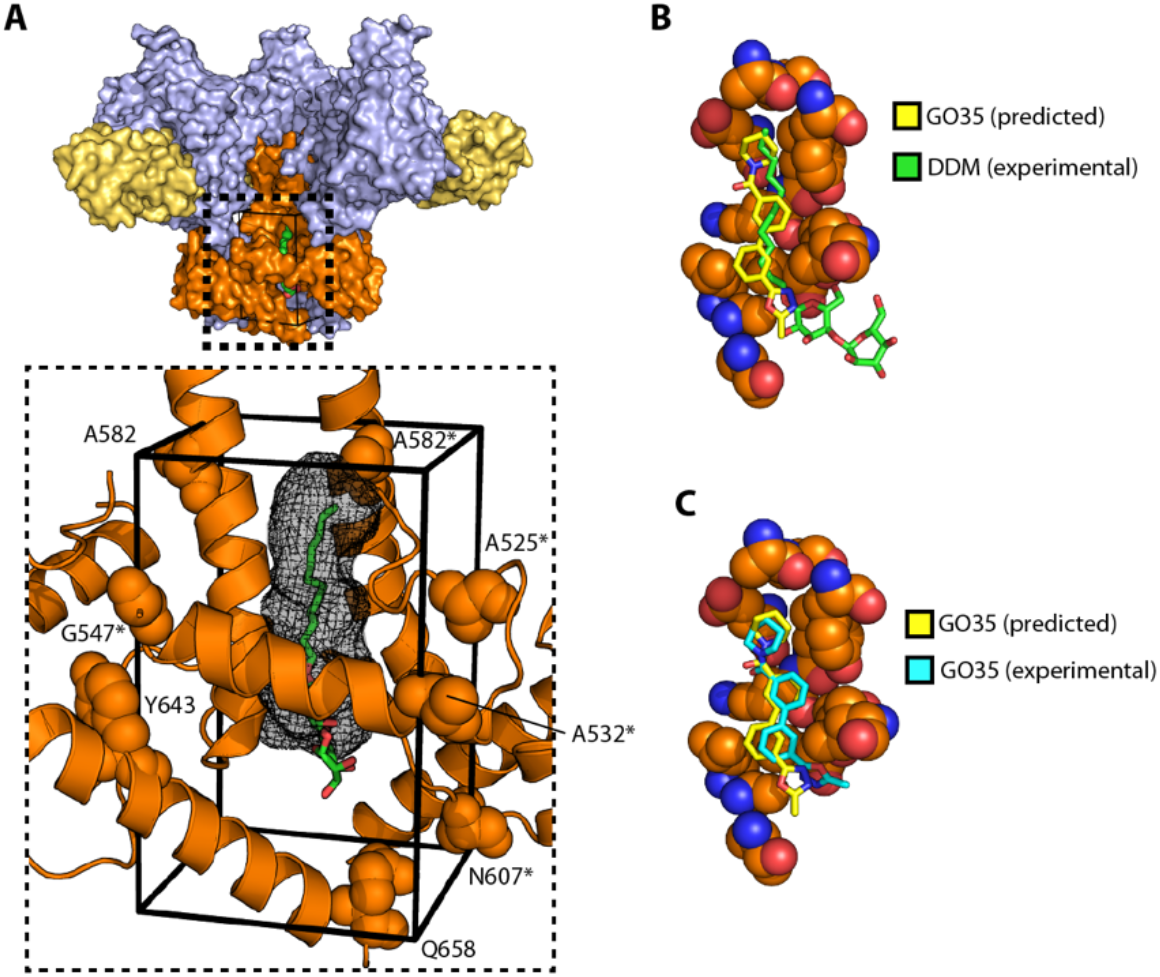
The DDM pocket as a template for *in silico* drug screening. (**A**) Location of the docking box with respect to the coordinates of DDM (*green* sticks), and the predicted AutoSite ligand binding site (black mesh); residues delimiting the box are shown as *orange* spheres; (**B**) Experimental coordinates of GO35 (*yellow* sticks) and DDM (*green* sticks) in the binding site (residues within 5 Å from any GO35 atoms as *orange* spheres; I519, P522 and A541 omitted for sake of clarity;); (C) Experimental (*yellow* sticks) and docking predicted (*cyan* sticks) coordinates of GO35 in the binding site (residues within 5 Å from any GO35 atoms as *orange* spheres; I519, P522 and A541 omitted for sake of clarity).

We virtually docked a library of ~300k compounds in the pocket and applied several filters to prioritize 500 results for visual inspection. We tested the robustness of the docking protocol by repeating the VS on the three pockets in the asymmetric reconstruction, and the results highlight two major aspects; on one hand, despite structural variations resulting from the asymmetric conformational changes, the conserved topology of the pocket is sufficient to reproduce the overall ranking of the hits selected for testing using the initial C3 symmetric structure; on the other, the loss of interactions with FP in the more disordered conformation reduces the magnitude and range of the docking scores (**Fig. S2, A to C**), possibly affecting the discriminatory power of the structure in separating binders from non-binders. Therefore, we focused on compounds displaying significant overlap with DDM density from the docking results into the C3 symmetric structure and purchased 59 for further investigation.

### A candidate molecule binds near the FP and interacts with conserved F522

To quickly assess potential binding of a candidate molecule to Env SOSIP, we hypothesized that a binding event might be inferred from a change in thermostability of the protein. Previously, we showed that analogs of known fusion-inhibitor BMS-626529, which binds to gp120, significantly increased the melting temperature of SOSIP trimers(*21*). We screened our compounds against 6 different Env SOSIP constructs (representing subtypes A, B or C) using differential scanning fluorimetry (DSF) by incubating a molar excess of small molecule with a complex of SOSIP and sCD4, and measuring the relative change in the thermal transition midpoint temperature (ΔT_m_) from a control containing the protein complex (SOSIP+sCD4) in 1% DMSO (**Fig. 3A, and table S2**). Candidate small molecules were chosen if they met both of the following criteria: 1) a ΔT_m_ value equal or greater than ±1.0°C, and 2) reactivity against at least 2 different Env genotypes. 8 of the compounds had intrinsic fluorescence that interfered with the method, and from the remaining 51 candidates, 5 were selected by the above criteria (**Fig. S2, D and E**). When compared to assays in which sCD4 was excluded, one small molecule, GO35, stood out as the change in T_m_ of 3 Env trimers was only observed in the presence of CD4, suggesting that the ligand is specific for the CD4-bound conformation (**Fig. 3B, fig. S2D, and table S2**). This small molecule decreased the thermostability of the CD4-bound complex by about 3°C, which we inferred as a possible conformational change, and it was chosen as the first candidate for structural studies.

**Fig. 3.**
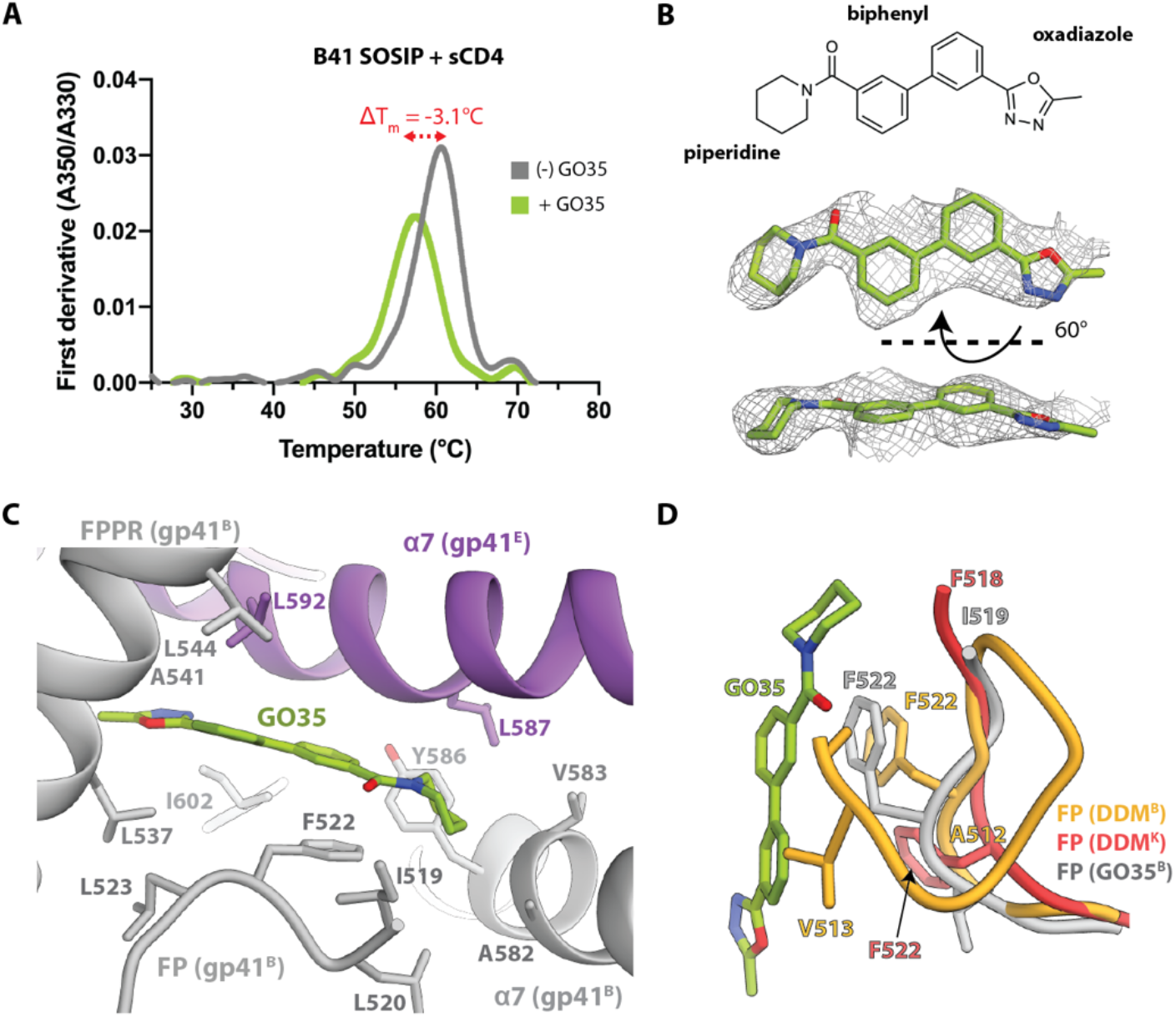
GO35 affects the thermostability of the receptor-bound Env complex and binds near the fusion peptide. (**A**) Differential scanning fluorimetry first derivate curves of CD4-bound B41 SOSIP in the presence or absence of GO35. (**B**) Chemical structure of GO35 and atomic coordinates and corresponding EM density of modeled GO35. (**C**) GO35 binding pocket and interaction with the fusion peptide. (**D**) Comparison of fusion peptides from DDM-bound and GO35-bound structures reveals that GO35 requires a different FP conformation to avoid steric clash.

It was imperative that a different detergent was used for cryo-EM vitrification to decrease chances of cross-competition of DDM with candidate small molecules. DSF analysis showed that lauryl maltose neopentyl glycol (LMNG) did not have a major effect on protein stability and has twice the mass of DDM, making it unlikely to fit into the binding pocket (**Fig. S3, A and B, and table S2**). As a control, we solved a ~3.7 Å cryo-EM structure of CD4- and 17b-bound B41 SOSIP frozen in the presence of LMNG (**Fig. S2, F and G, and table S1**). Not only did we not see any additional density in the FP pocket that could account for detergent, but the FP itself was less ordered, similar to that of the partially-bound pocket in the DDM reconstruction (**Fig. S2H**), supporting our hypothesis that the presence of ligands affects the local stability of the FP.

Using single particle cryo-EM, we next solved a ~3.5 Å C3 symmetric reconstruction of a complex of B41 SOSIP, sCD4 and 17b Fab that was incubated with GO35 (**Fig. S3, C to E, and table S1**). The N-terminal portion of the fusion peptide is disordered until residue I519 (**Fig. 3C**). Density for the entire GO35 molecule is sandwiched between the FPPR helix and FP of gp41^B^ and HR1 helix of gp41^E^ (**Fig. 3A,C**). One of the biphenyl rings stacks against the side chain of conserved F522 (>98% of all HIV sequences) of the fusion peptide (**Fig. 3C**), and comprises the most extensive interaction with Env. The second ring of biphenyl is stabilized by a hydrophobic local environment consisting of L537^B^, A541^B^, L544^B^, L592^B,E^, and I602^B^. The piperidine ring of GO35 is located deepest in the pocket, although the density supports only weak hydrophobic interactions with the environment of I519^B^, L520^B^, Y586^B^, A582^B^, V583^B^, and L587^E^ (**Fig. 3C**). On the other side of the central biphenyl is the oxadiazole ring near the entrance to the pocket, and it may form hydrogen bonds with the peptide backbone of the FPPR (**Fig. 3C**).

Remarkably, the experimental binding mode of GO35 overlaps substantially with the position of DDM in the model used as a reference (**Fig. 2B**), and with minimal deviation from the predicted binding mode (RMSD 1.7 Å) (**Fig. 2C**). When compared to the asymmetric DDM-containing model, the FP of the GO35-bound model is resolved only from I519, similar to the more disordered conformation seen in DDM-bound gp41 chain K, which is adjacent to a partially occupied pocket (**Fig. 3D**). However, residues 519-525 align best with the equivalent region of DDM-bound gp41 chain B (adjacent to a fully resolved DDM molecule), particularly the side chains of F522. The full fusion peptide from the DDM-bound conformation would potentially clash with GO35 as the FP folds back on itself, bringing A512 and V513 very close to the biphenyl core of the small molecule (**Fig. 3D**). This suggests that the binding of GO35 biases the FP toward the more disordered state and may force the trimer into a less stable conformation as trimer dissociation was apparent in the cryo-EM 2D class averages, with an estimated ~60% of the selected particles classified as individual protomers (**Fig. S3F**).

### A second small molecule is capable of low micromolar inhibition

We next used a TZM-bl cell assay to measure whether GO35 neutralizes HIV (**Fig. 4, A and B**). Molecules were tested for cytotoxicity up to 60 μM and GO35 did not have a measurable effect, so dilutions of the compound were tested in neutralization assays up to half of this value (30 μM) (**Fig. S4A**). Despite the experimentally-determined binding of GO35 to B41, neutralization was not seen against this virus nor 13 other genetically diverse HIV-1 viruses, motivating us to further examine other small molecule hits from the virtual screen. (**Fig. 4A**).

**Fig. 4.**
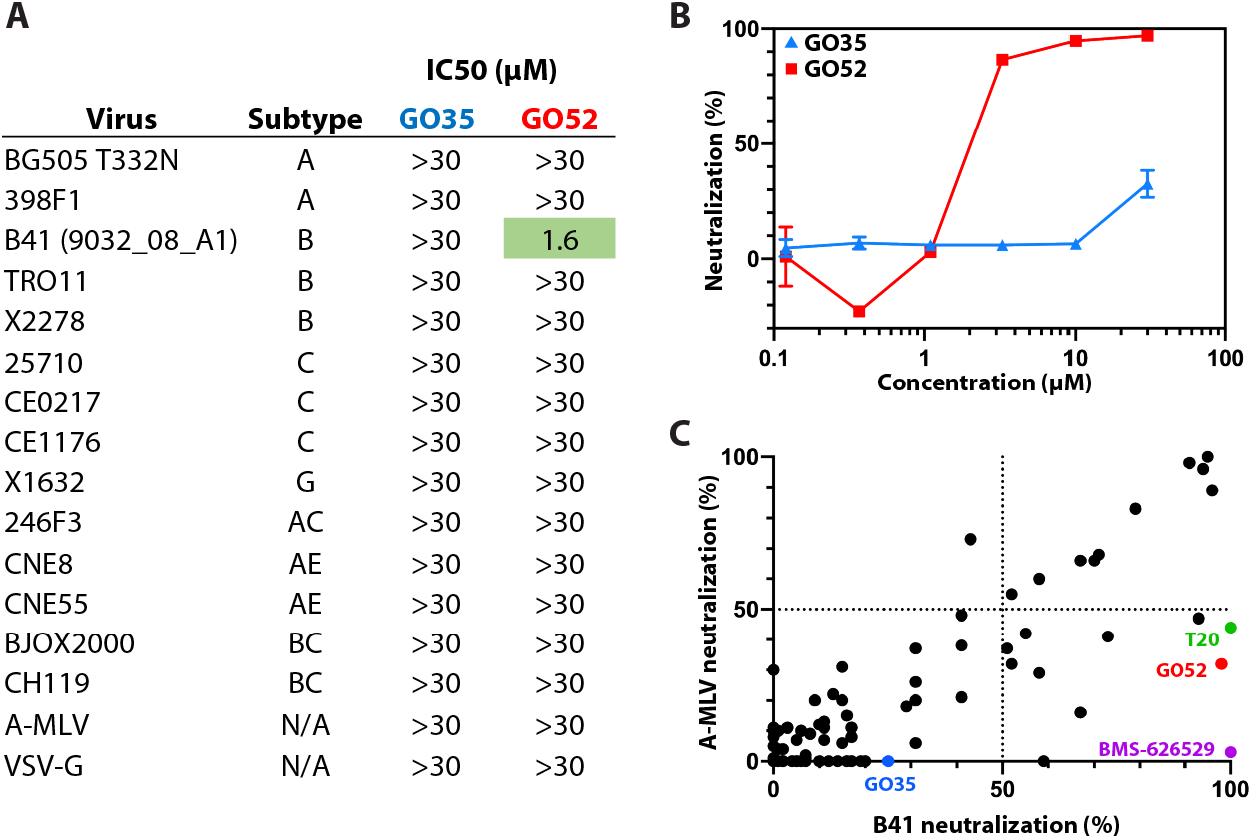
Screening for other hits using HIV-1 neutralization assays. (**A**) Neutralization profiles of GO35 and GO52 against 14 HIV-1 and 2 control viruses. Assays performed as duplicates (*N=2*) and IC50 determined by fitting an asymmetric sigmoidal five-parameter dose-response curve (R^2^=0.9757 for fit of GO52 against B41 virus). (**B**) TZM-bl neutralization curves of GO35 and GO52 against B41 pseudotyped virus. Assay performed as duplicates (*N=2*) and error bars represent standard deviation. (**C**) Neutralization activity of 80 small molecules against B41 HIV-1 and A-MLV viruses. All molecules tested at 30 μM final concentration. Assay performed as duplicates (*N=2*) and mean value is plotted. BMS-626529 and T20 (enfuvirtide) are known HIV-1 fusion inhibitors included as controls.

Because DSF should not be expected to pick up all binding events, nor necessarily correlate with neutralization, we screened ~80 compounds (including analogs of GO35, and known fusion inhibitors T-20 [enfuvirtide] and BMS-626529 as positive controls) using the TZM-bl neutralization assay against B41 and A-MLV (negative control for HIV specificity). This screen revealed a few promising HIV specific hits, including GO52 (**Figs. 4C and 5A**). Further assays measured an average IC50 of 1.6 μM for GO52 against B41, although the small molecule was not able to neutralize other viruses in the 12-member global panel (**Fig. 4, A and B**). At high concentration (30 μM) some neutralization against the negative control A-MLV was measured (**Fig. 4C**). Interestingly, the T-20 control also showed some non-specific effects against A-MLV, but this non-specificity was not seen for BMS-626529 (**Fig. 4C**). Due to its measured inhibition and relative specificity towards HIV-1, we next investigated GO52 further by cryo-EM.

**Fig. 5.**
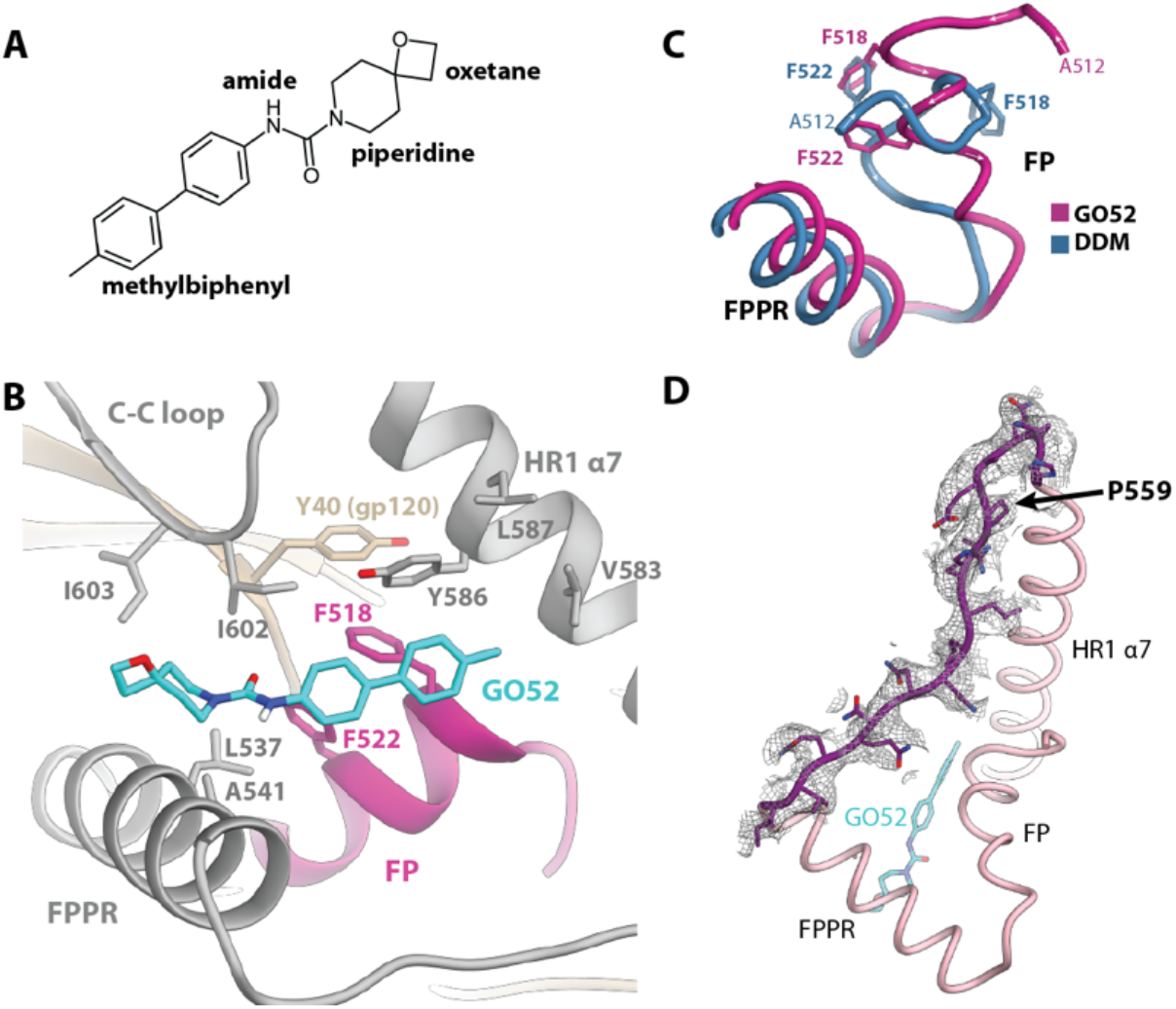
GO52 stabilizes a new conformation of the FP and surrounding regions. (**A**) Chemical structure of GO52. (**B**) GO52 binding pocket and interactions with surrounding peptide. (**C**) Comparison of fusion peptides from DDM-bound and GO52-bound structures a large rearrangement in which F518 (GO52-bound) takes the place of F522 (DDM-bound). (**D**) Model and EM map for residues 548-564 of HR1, a region that is typically disordered in published structures.

### GO52 binds the trimer base via aromatic interactions with gp41

During our previous cryo-EM attempts, we noticed that the complexes, particularly the trimers, had a tendency to dissociate over time in the presence of small molecules (**Fig. S3F**). Presumably, this destabilization occurs from the presence of the solvent, or even the trapping of an energetically unfavorable state of the FP. To circumvent this, we used glutaraldehyde to crosslink the B41-CD4-17b complex to stabilize it prior to small molecule addition. Indeed, fewer dissociated protomer particles were seen in the frozen samples, with an estimated 28% of particles resembling dissociated protomers compared to ~60% in the GO35 sample (**Fig. S5A**). Ultimately, a 3.6 Å cryo-EM reconstruction revealed that GO52 binds in the predicted pocket, and major interactions involve conserved Y586 and F522, and a rarely occurring (1.5% of sequences) F518 that is found in B41 Env (**Fig. 5B, fig. S5B, and table S1**). The FP is reconfigured compared to the DDM-bound complex such that the side chain of F518 supplants the F522 side chain from the DDM-bound complex. F522 now becomes a secondary contact to GO52 and is trapped between F518 and L537 (of FPPR) (**Fig. 5C**). F518 forms a cluster of five aromatic rings (F518, F522 and Y586 of gp41, Y40 of gp120, and the methylbiphenyl group of GO52) (**Fig. 5B**).

The C3-symmetric map suggests dynamic movement of the FP centered on F522 that is not simply a difference in rotamers, so we generated an asymmetric reconstruction of the same dataset (~4.0 Å resolution) to investigate further (**Fig. S5C, and table S1**). In all three protomers, the phenyl group of F522 appears centered between the side chains of L537 and F518 resulting in hydrophobic and π-π stacking interactions and the cluster of 5 aromatics is preserved (**Fig. S5D**). The extra density near F522 in at least one protomer appears to be from hydrophobic interactions between P43 (of gp120) and the α, β and γ carbon atoms of F522, while in the other two protomers, P43 interacts with either the side chains of L523 or A526 (**Fig. S5F**). Portions of the FP in all 3 protomers appears to form an α-helix (**Fig. S5D**).

With few exceptions, a portion of gp41 HR1 (residues 548 to 660, HXB2 numbering) is disordered in published Env structures, whether in the receptor-free or receptor-bound pre-fusion states. Sometimes the binding of a gp120-gp41 interface antibody (PGT151) confers more stability to this region, or the engineering of a more stable crystal lattice (*6, 11*). This region is also disordered in our DDM- or GO35-bound structures. Interestingly, the entire region is ordered in the GO52-bound structure and can be fully modeled (**Fig. 5D**). While we cannot exclude the possibility that glutaraldehyde cross-linking is responsible for this observation, the HR1 region and surrounding residues do not contain lysine residues that are most likely to be modified by glutaraldehyde (**Fig. S5E**) Furthermore, the region is also disordered in a published BG505 Env trimer that had been crosslinked with glutaraldehyde (*22*). It is possible that increased local order is therefore a result of GO52 binding.

While GO52 causes a global destabilization of the trimer it does induce a relatively homogenous and stable association between the FP, FPPR, α7 HR1 helix and C-C loop of gp41, and C1 gp120, with possible allosteric effects on gp120 C5 (via C1 stabilization). In comparison, the C1 model of DDM-bound B41-CD4-17b exhibits greater asymmetry in this region, with F518 not playing a stabilizing role, and both F518 and F522 dramatically translocating in one of the protomers relative to the other two (**Fig. 3D**). Furthermore, the FP is not stabilized into an α-helix in any of the protomers.

### Differences exist between receptor-induced Env rearrangements across genotypes

The specificity of GO52 against B41 prompted us to investigate whether this was due to simple amino acid variation or a larger difference of pocket accessibility across genotypes. As mentioned above, a phenylalanine residue at position 518 of the fusion peptide is rare (1.5%; 91/5923 sequences in the Los Alamos Database). Perhaps this residue is key for neutralization so we obtained two viruses (clade G X1254.c3 and clade AG T251-18) that naturally have F518 and only deviate from B41 by single amino acid substitutions in the FP (**Fig. 6A**). Despite the similarity, GO52 demonstrated no neutralization activity against these pseudotyped viruses (**Fig. 6A**).

**Fig. 6.**
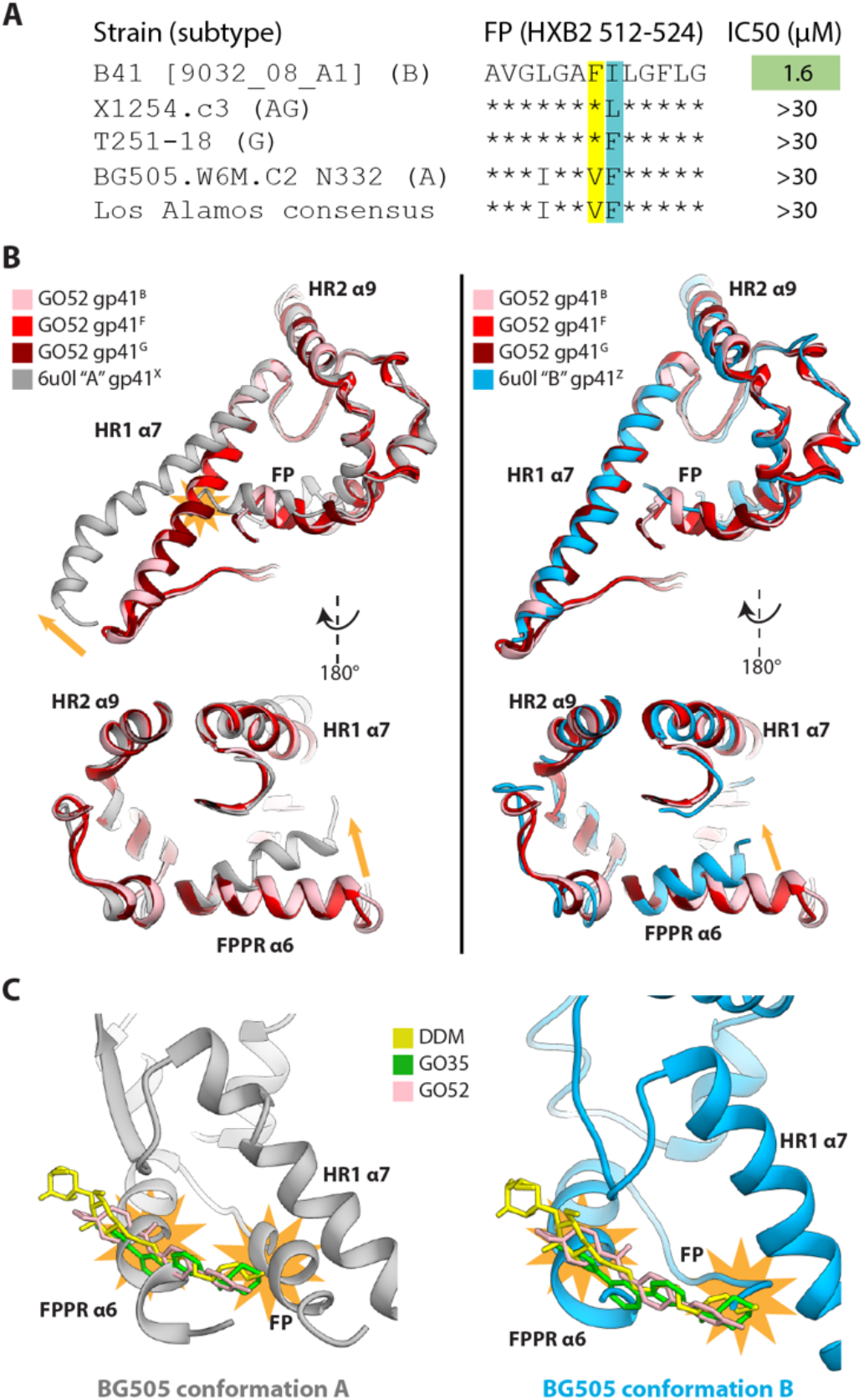
The B41 small molecule binding pocket is not amenable with published BG505 structures. (**A**) Sequence alignment of the fusion peptide (HXB2 512-524), relative to B41, of two HIV-1 sequences that also contain a phenylalanine at position 518, in comparison to BG505 and the Los Alamos consensus sequence. IC50 determined as in **Fig. 4**. (**B**) Overlay of CD4- and GO52-bound B41 (asymmetric) and CD4-bound BG505 conformation “A” (*left*) and conformation “B” (*right*) gp41 chains. Clashes denoted as orange stars, relative movements of BG505 with respect to B41 show as orange arrows. (**C**) Overlay of ligands from CD4-bound B41 with CD4-bound BG505 conformation “A” (*left*) and conformation “B” (*right*) gp41 chains. Clashes denoted as orange stars.

We next hypothesized that the binding pocket may differ across genotypes and compared our CD4-bound B41 Env structures to recently published CD4-bound BG505 Env structures (that contain E51 as the co-receptor mimic antibody, instead of the 17b antibody in our structures)(*18*). A notable feature of the BG505 complex is that two different cryo-EM classes suggest a high degree of asymmetry between protomers. When comparing gp41 protomers from the BG505 structures to B41, a shift in the relative angle of the HR1 α7-helix is apparent, in which the BG505 “conformation A protomer” helix pivots further away from the FP pocket relative to B41 (**Fig. 6B**). The N-terminal portion of the BG505 FP clashes with the α7-helix orientation of B41. In addition, the FPPR helix (α6) is shifted more into the FP pocket in the BG505 structure and constricts the B41-derived drug pocket (**Fig. 6B**). The repositioned FPPR and FP clash with the modeled B41 ligands (DDM, GO35, GO52) (**Fig. 6C**). The “conformation B protomer” has better alignment to B41, with high overlap of α7- and α9-helices, but also has the shifted FPPR helix (**Fig. 6B**).

Our binding pocket derived from a B41 model is different than available models of BG505. The BG505 FP sequence is identical to the overall consensus sequence based on 5,923 Los Alamos HIV Database entries (**Fig. 6A**). If the FP dictates the gp41 conformational changes then it is possible that other genotypes have a CD4-bound structure more similar to BG505 than B41, explaining the observed specificity of GO52. In fact, when we screen our top hits derived from B41 docking against BG505 and correlate it with an A-MLV negative control, we find that none of our compounds show neutralization specificity against BG505 (**Fig. S4**). However, even the CD4-bound BG505 data suggest conformational heterogeneity in this region. Although the overall maps are of high quality and resolution (3.3 and 3.5 Å overall, with no local resolutions reported that are higher than the global FSC), a comparison of the density of FPPR and surrounding regions compared to our DDM-bound models suggests that in BG505 this region is more flexible as evidenced by more disorder in the maps (**Fig. S6**).

## Discussion

The serendipitous discovery of a detergent molecule residing in a receptor-binding induced pocket provided the framework for investigating whether this could be exploited for HIV fusion inhibitor development. Our initial hit based on binding alone, GO35, influenced the conformation of the FP by a π-π stacking interaction with a conserved phenylalanine residue, but did not neutralize. GO52 showed inhibitory activity specific to the parental virus of the docking model Env, B41. Comparisons to BG505 suggest that the receptor-bound conformational state varies across genotypes. It is possible that B41 naturally forms the binding pocket more often and homogenously across the 3 binding sites, and is therefore more available to binding the small molecule. The conserved F522 residue might play a major role in regulating this state of the Env trimer. We have described before how CD4-binding site antibody b12 mimics the receptor-induced conformational changes in gp41, including those of the FP, while the antibody does not bind to or neutralize BG505 virus (*7*).

We initially used a cryo-EM-derived model as a target for docking and virtual screenings, generated top candidates that were screened using cell-based and biophysical assays, and then solved two more cryo-EM structures, each containing unambiguous evidence of small molecule binding (**Fig. S7**).

Although our efforts did not reveal molecules capable of neutralizing multiple HIV-1 strains, we did, however, succeed in finding an inhibitor that binds the specific pocket of our search model. Furthermore, the initial model has the conserved F522 in the binding pocket while the rare F518 was outside of it. Thus, the resulting conformational change induced and/or trapped by GO52 came as a surprise. Collectively, the results of this study not only support the viability of cryo-EM to provide data for atomistic modeling of potential drugs, which continues to be corroborated for other important drug targets (*23–25*), but they also constitute a success story of the use of molecular docking with cryo-EM structures. In fact, the approach described here has worked successfully for finding new molecules that bind to a transient target, such as an HIV pre-fusion intermediate, while capturing multiple conformational ensembles. Our results demonstrate the success of the *in silico* screens for being selective against our desired target. Future efforts will focus on elucidating the structure of this binding pocket using another Env genotype, one that has a more conserved fusion peptide sequence.

## Materials and Methods

### Experimental design

#### Binding site analysis

AutoSite(*26*) with default settings was used to analyze the initial C3 env model and identify the optimal ideal ligand volume for the DDM site. The program predicted correctly the three DDM sites (**Fig. 2**), with optimal volumes overlapping with the hydrophobic tail of the detergent and the first glucose ring.

#### Docking

The ChemBridge ligand library (1.3M compounds [https://www.chembridge.com/] (accessed February 2019)]) was downloaded from ZINC(*27*) [http://zinc.docking.org/ (accessed August 2016)], and filtered to obtain the 90% diversity set (301k ligands). Ligands were then prepared according to the standard AutoDock protocol(*28*). The refined model was used to extract the coordinates used for the dockings, which included two adjacent monomers of gp-41 (chain B and E), and one monomer of gp-140 (chain A). Following the standard AutoDock protocol(*28*), the structure was prepared by removing all non-standard amino acids, as well as glycans (no glycosylation sites were included in the docking box). Explicit hydrogens were added with Reduce(*29*). AutoDock Vina v.1.1(*30*) was used to perform docking calculations using a docking box centered at coordinates x=158.694, y=161.962, z=133.764, and sized 20.62 x 20.62 x 33.75 Å. Results were filtered and analyzed using AutoDock Raccoon2(*28*), discarding compounds with predicted score of less than −13.6, and ligand efficiency of −0.26 or worse. Through visual inspection, 59 compounds were selected and purchased. The ZINC IDs for all 59 compounds can be found in **Table S2**.

#### Small Molecule Stocks

Small molecules were purchased from ChemBridge (San Diego, CA). Master stocks of the small molecules (with an approximate molecular weight of ~330 Daltons) were created by dissolving in 100% DMSO at a concentration of 20 mg/ml (~60,000 μM). All stocks were stored at −20°C. In addition, BMS-626529 (APExBIO; dissolved to 20 mg/mL [~42,000 μM] in 100% DMSO) and known fusion inhibitor, T20 (enfuvirtide, dissolved in PBS to 5 mg/mL [~1,110 μM]; Sigma Aldrich) were purchased to serve as positive controls for neutralization assays.

#### Protein Expression

AMC011 v4.2 SOSIP.664, BG505 SOSIP.664, B41 SOSIP.664, CZA97 SOSIP.664, DU422 SOSIP.664, and JRFL SOSIP.664 trimers were transiently transfected with Furin in HEK-293F cells (Invitrogen) and purified with in-house made PGT145 immuno-affinity columns, followed by size exclusion chromatography on an AKTA Pure paired with a HiLoad 16/600 Superdex 200 prep grade column (General Electric Healthcare) using methods previously described(*31, 32*). 17b Fab was expressed in ExpiCHO cells (Invitrogen), purified using a 1 ml Thermo Capture Select column (Thermo Fisher), and eluted with 0.1 M sodium acetate pH 3.5. Fractions of interest were pooled, concentrated, and buffer exchanged into TBS, with a 10 kDa concentrator (Millipore Sigma). Soluble CD4 was transiently transfected in ExpiCHO using ExpiFectamine, expressed for 14 days using the Max Titer Protocol, and purified as described previously(*7*).

#### Nano-Differential Scanning Fluorimetry (DSF)

Thermostability tests were performed using a nano-DSF Prometheus NT.48 instrument and standard grade capillaries (Nano-Temper Technologies). In each instance, the samples at ~0.2 mg/ml were subjected to a temperature variance of 20°C to 95°C, using a thermal ramp of 1°C per minute. Values reported correspond to the inflection point calculated within the PR.ThermControl software (Nano-Temper Technologies).

#### Small molecule screen

A panel of 59 small molecule candidates were screened for intrinsic fluorescence. To do this, 1:100 dilutions of the small molecule master stocks were made with 1X TBS pH 7.4. The candidate molecules were resuspended in DMSO, and in all cases diluted prior to DSF experiments such that the final DMSO concentration was 1% (v/v), which was found to have a negligible impact on the melting temperature of SOSIP trimers.

#### SOSIP.664 trimers in complex with sCD4, and small molecules

10 μL of AMC011 v4.2 SOSIP.664, BG505 SOSIP.664, B41 SOSIP.664, CZA97 v3 SOSIP.664, DU422 SOSIP.664, and JRFL SOSIP.664 at 1 mg/ml were complexed with sCD4 at 1 mg/ml and each small molecule that did not show intrinsic fluorescence (GO17, 28, 30, 32, 40, 44, 47, 58 were excluded). Final concentrations of SOSIP, sCD4, and small molecules were 0.94 μM, 5 μM, and 400 μM, respectively.

#### Cytotoxicity Assays

A 3-fold dilution series of small molecules GO35 and GO52 was prepared in 100% DMSO ranging from 9,000 to 4.12 μM. 1 μl of each small molecule was added to an individual well in a black 96-well flat bottom plate, followed by 99 μL of TZM-bl cells at 0.1 million cells/ml containing 10 μg/ml DEAE-Dextran (Sigma Aldrich). Final concentrations of the small molecules were 90, 30, 10, 3.3, 1.1, 0.37, 0.12, and 0.04 μM in 1% (v/v) DMSO. Each condition was set up as duplicates. 4 reference wells containing 1% (v/v) DMSO and cells were also included, along with 4 control wells containing 1% (v/v) DMSO in DMEM media. After 2 days of incubation, 100 μL of CellTiter-Glo 2.0 Reagent (Promega) was added to each well. Plates were transferred to a Biotek Synergy H1 Microplate Reader, subjected to orbital shaking for 2 minutes followed by a 10 minute delay before luminescence was measured. Viability of the cells was calculated from values generated in the Gen5 software (Molecular Devices) by normalizing using the mean values of the DMSO and cell only wells (100% viability) and the DMSO and media only wells (0% viability). Normalization and generation of plots was performed using GraphPad Prism version 8.

#### Pseudovirus Neutralization assays

Plasmid DNA for pseudotyped viruses, HEK293T and TZM-bl cells were obtained from the NIH AIDS Reagent Program and as a gifts from Dr. John Moore (Weill Cornell Medical College) and Mark Louder (NIH Vaccine Research Center). All pseudotyped HIV-1 Env viruses used in this study were produced by co-transfection with the NL4-3 plasmid in HEK293T cells following previously described(*33*) protocols and the standardized protocols of Dr. David Montefiori (https://www.hiv.lanl.gov/content/nab-reference-strains/html/home.htm). Small molecule neutralization assays were performed according to a standard TZM-bl protocol(*33*) with pseudotyped HIV-1 viruses.

Small molecules dissolved in DMSO were diluted 1:100 such that the final DMSO concentration was 1% (v/v), and final small molecule concentrations ranged from 0.01-30 μM. Each condition was tested in duplicate. Both virus and cell-free control plates contained 1% (v/v) DMSO. Small molecules were incubated with pseudotyped viruses for 1 hour at 37°C prior to the addition of TZM-bl cells. The calculated amount of DEAE-Dextran (Sigma Aldrich) used in the assays was 10 μg/ml. Unused wells around the perimeter of the plate were filled with DPBS to minimize evaporation of the experimental wells. After 48 hours, the Britelite Plus Reporter Gene Assay System (PerkinElmer) was added and plates were transferred to a Biotek Synergy H1 Microplate Reader and recorded using the luminescence filter and Gen5 software (Molecular Devices). Normalization was performed by calculating the mean values for reference wells containing virus, cells and 1% DMSO (0% neutralization) and the virus-free wells containing DMSO and cells (100% neutralization). Normalization, titration curves and IC50 calculations were performed using the GraphPad Prism software (version 8) by fitting an asymmetric sigmoidal five-parameter dose-response curve.

#### Negative stain electron microscopy

CZA97 SOSIP.664 trimers were incubated overnight at room temperature with a six-fold molar ratio of soluble CD4 and a ~400-fold excess of small molecules GO20, 23, 34, 35, and 57. The following day, complexes were diluted to ~0.01 mg/ml with 1X TBS pH 7.4, deposited on glow discharged copper mesh grids (Electron Microscopy Sciences), and negatively stained with 2% uranyl formate. A 120 keV FEI Tecnai Spirit with a TIETZ 4K x 4K camera was used to collect data, facilitated by the Leginon software(*34*). Micrographs were stored and processed within the Appion database(*35*). Complexes were picked using DogPicker(*17*), stacked with a box size of 160 pixels, and 2D classification was performed with iterative multivariate statistical analysis/multireference alignment (MSA/MRA)(*36*). Any obvious particle contaminants were removed from the classification.

#### Cryo-Electron Microscopy sample preparation

Preparation of B41+CD4+17b frozen with DDM was previously described(*7*). For complexes of B41+CD4+17b with small molecule GO35 or GO52, ~400 μg of B41 SOSIP.664 were incubated overnight at room temperature with sCD4 and 17b Fab, both at an approximate six-fold molar excess to SOSIP. The mixture was size-exclusion purified the following day with a HiLoad Superdex 200 pg column (GE Healthcare), and appropriate fractions were concentrated to ~50 μL with a 100 kDa molecular weight concentrator (Amicon Ultra, Millipore). For the sample intended for incubation with GO52, an equal volume of 15 mM glutaraldehyde (Thermo Fisher Scientific) was added and incubated for 30 min, and the reaction was quenched by adding 1 M Tris (Thermo Fisher Scientific) to a final concentration of 0.1 M. Size-exclusion chromatography was performed a second time on the cross-linked sample. Final concentrations of the complexes were ~5 mg/ml (GO35) and ~1.2 mg/ml (GO52). Either small molecule GO35 or GO52 was diluted 1:100 into the complex (~600 μM or 150 μM final concentrations for GO35 or GO52, respectively; ~26x molar excess) and incubated for less than 30 minutes. To aid in particle orientation distribution, 0.5 μL of lauryl maltose-neopentyl glycol (LMNG, Anatrace) at 0.04 mM and 3.5 μL of either complex were briefly incubated prior to deposition onto Solarus plasma cleaned (Argon/Oxygen) Quantifoil 1.2/1.3-200 mesh or C-flat 2/2-400 mesh copper grids. Samples were plunge-frozen with a Thermo Fisher Vitrobot Mark IV at 10°C, 100% humidity, 10 second wait time, and a blot force of 0. The Quantifoil 1.2/1.3-200 mesh grid required an 8 s blot time, while the C-flat 2/2-400 mesh grid was blotted for 4.5 s.

#### Cryo-EM data collection and processing

Relevant map and model statistics are summarized in **Table S1**. All data were collected on a Titan Krios (Thermo Fisher) operating at 300 keV or Talos Arctica (Thermo Fisher) operating at 200 keV, each equipped with a Gatan K2 Summit camera. Raw micrographs were aligned and dose weighted using MotionCorr2(*37*). These micrographs were then imported into cryoSPARC version 2(*16*). CTF was estimated using GCTF(*38*), and micrographs with a CTF fit resolution above 5 Å were discarded. Particles were picked using the template picker and were subsequently extracted with a box size of 288 pixels. Subsequent processing was continued in either cryoSPARC version 2 (*16*) (DDM, LMNG, and GO35 datasets) or particles were exported to Relion 3.0 (*15*) (GO52 dataset). After numerous rounds of 2D classifications and 3D sorting, the final particles were subjected to non-uniform refinement (cryoSPARC version 2) or 3D auto-refine, CTF refinement and post processing (Relion 3.0).

#### Model Building

PDB 5VN3 (B41 SOSIP in complex with sCD4 and 17b) was used as an initial model for all datasets and fit into each respective map using UCSF Chimera(*39*). Due to masking and non-uniform refinement methods to improve resolution at the GO35-binding site, 17b Fab was excluded from the GO35- and GO52(C1)-bound models as the density for this region was of lower resolution. The coordinates for DDM were imported from the REFMAC5(*40*) dictionary in COOT(*41*), while the coordinates for GO35 and GO52 were generated using Phenix eLBOW(*42*) and the ZINC(*27*) SMILES string. Refinement was performed using Rosetta Relax(*43*) and models were validated using MolProbity(*44*) and EMRinger(*45*) included in the Phenix software suite(*46*).

### Statistical Analysis

Neutralization assay curves and IC50 determination were using GraphPad Prism (version 8.4.2) software. All statistical measures are clearly described in the figure legends and/or in the Materials and Methods.

## Acknowledgments

**General:** We thank J.C. Ducom and Lisa Dong at the Scripps HPC facility for the computational support, as well as Bill Anderson, Hannah Turner and Charles Bowman for help with electron microscopy data collection and processing. This is manuscript #29845 from The Scripps Research Institute.

## Author contributions

All authors conceived and designed the experiments, analyzed the data, and contributed edits to the manuscript. G.O., S.F. and A.B.W. wrote the manuscript. G.O., and J.L.T. performed cryoEM, DSF and neutralization experiments. D.S-M. and S.F. performed virtual screening experiments. J.L.T. expressed and purified protein reagents and pseudotyped viruses.

## Data and materials availability

CryoEM reconstructions and maps have been deposited to the Protein Data Bank and EM Data Bank under the accession numbers PDB 6OPN, 6OPO, 6OPP, 6OPQ, 6X5B, and 6X5C, and EMD-20150-20153, and EMD-22048-22049.

## Supplementary Materials

**Fig. S1.**
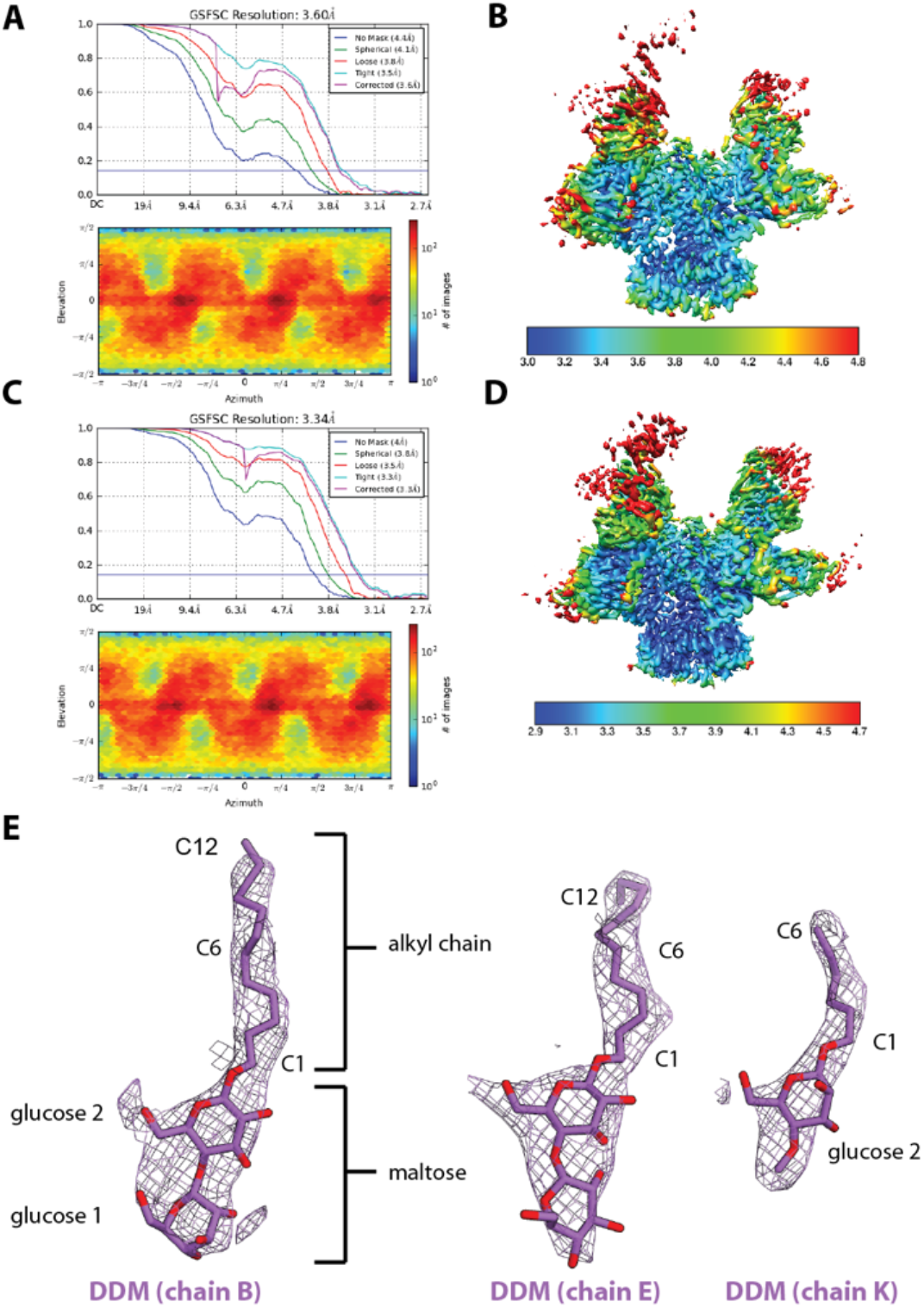
Cryo-EM maps and models of detergent-bound HIV Env in complex with receptor. (**A**) Fourier shell correlation (*top*) and distribution plot of particle orientations (*bottom*) and (**B**) local resolution estimates (colored by Å) of C1 reconstruction of B41-CD4-17b. (**C**) Fourier shell correlation (*top*) and distribution plot of particle orientations (*bottom*) and (**D**) local resolution estimates (colored by Å) of C3 reconstruction of B41-CD4-17b. (**E**) Density and modeled atoms for DDM in three protomers of the asymmetric reconstruction of B41-CD4-17b.

**Fig. S2.**
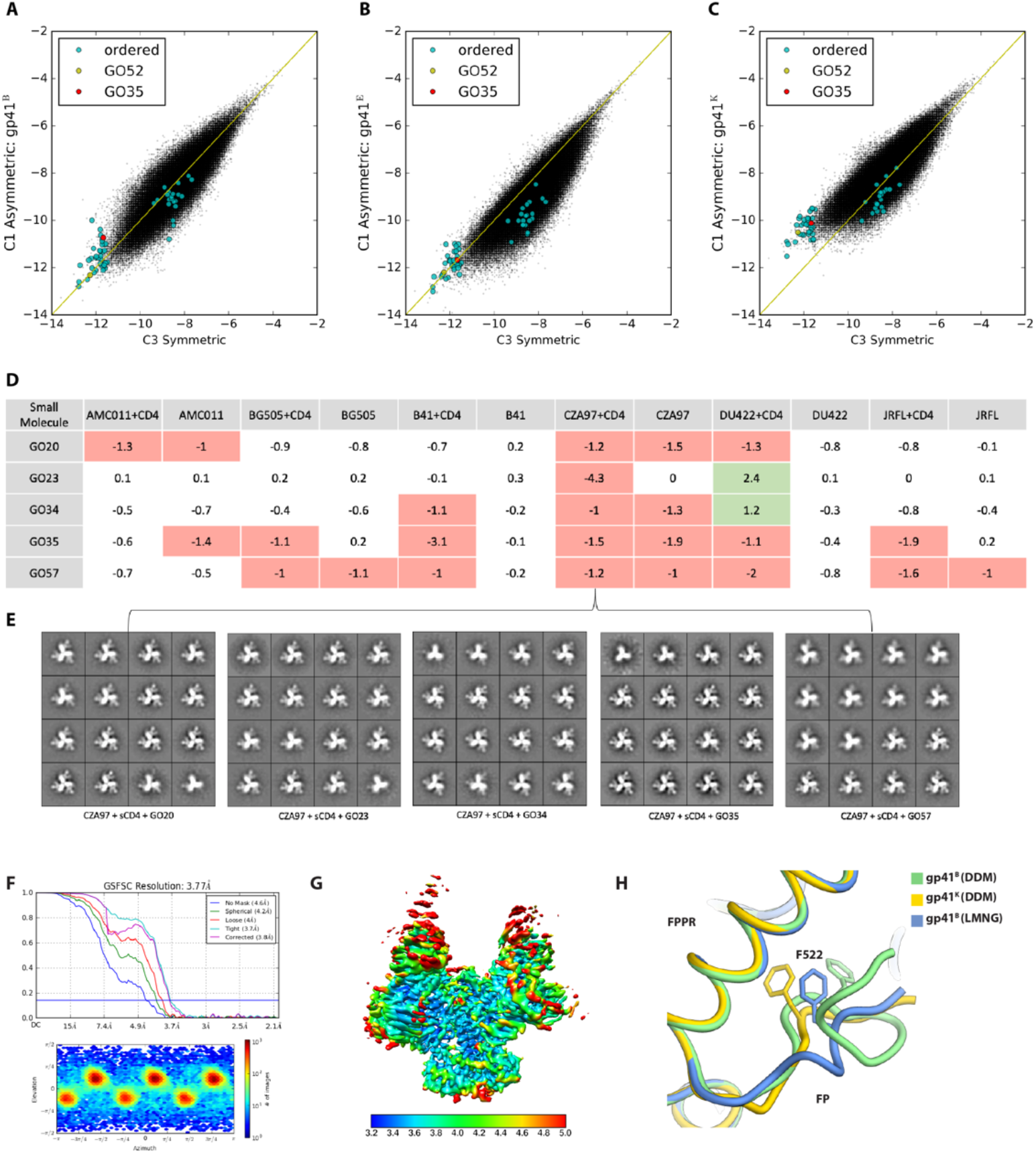
Selection of small molecule candidates by *in silico* screening and differential scanning fluorimetry. Plot of scores obtained in the C1 asymmetric structure (y-axis) against scores obtained in the C3 symmetric structure (x-axis). Each of the three panels corresponds to a different FP conformation as represented in **Fig. 1A**: (**A**) gp41^B^, (**B**) gp41^E^ and (**C**) gp41^K^. (**D**) Top 5 candidates based on a selection criteria of a ΔT_m_ value equal or greater than ±1.0°C against at least 2 different Env genotypes, in the presence or absence of sCD4. (**E**) Negative-stain 2D class averages of CZA97 SOSIP+sCD4 complexed with the top 5 candidates in (**D**) reveals that changes in stability are not due to trimer dissociation/denaturation. (**F**) Fourier shell correlation (*top*) and distribution plot of particle orientations (*bottom*) and (**G**) local resolution estimates (colored by Å) of C3 reconstruction of B41-CD4-17b frozen with LMNG. (**H**) Comparison of fusion peptides from B41-CD4-17b frozen with either DDM (asymmetric model) or LMNG.

**Fig. S3.**
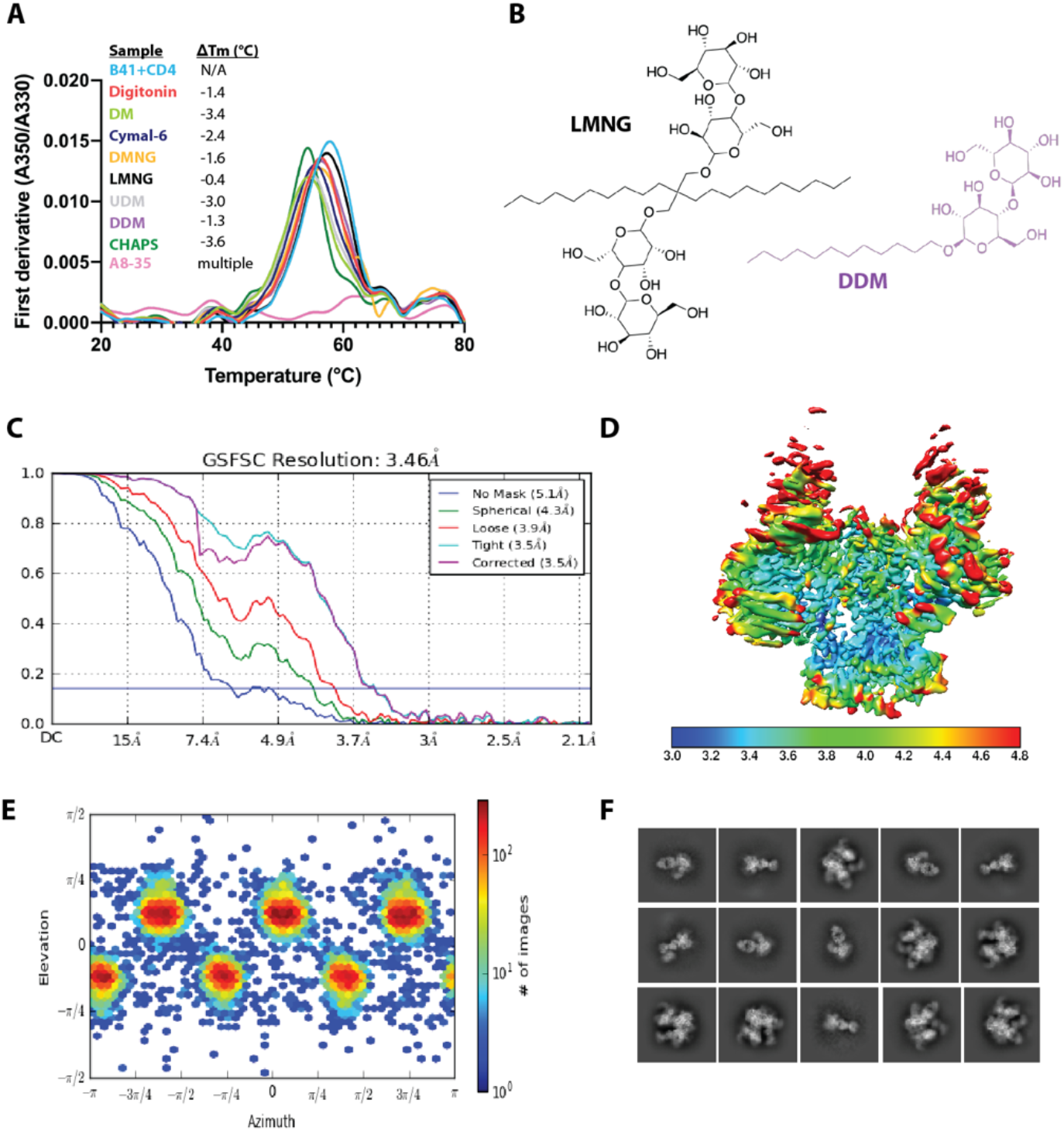
Targeting the DDM-pocket of gp41 with small molecules. (**A**) Change in T_m_ of B41 SOSIP in the presence of various detergents. (**B**) Comparison of DDM and LMNG structures. (**C**) Global Fourier shell correlation (FSC), (**D**) side view of the final 3D reconstruction colored by local resolution estimates (in Å), and (**E**) distribution plot of particle orientations of B41+CD4+17b+GO35. (**F**) Select 2D class averages of B41+CD4+17b+GO35.

**Fig. S4.**
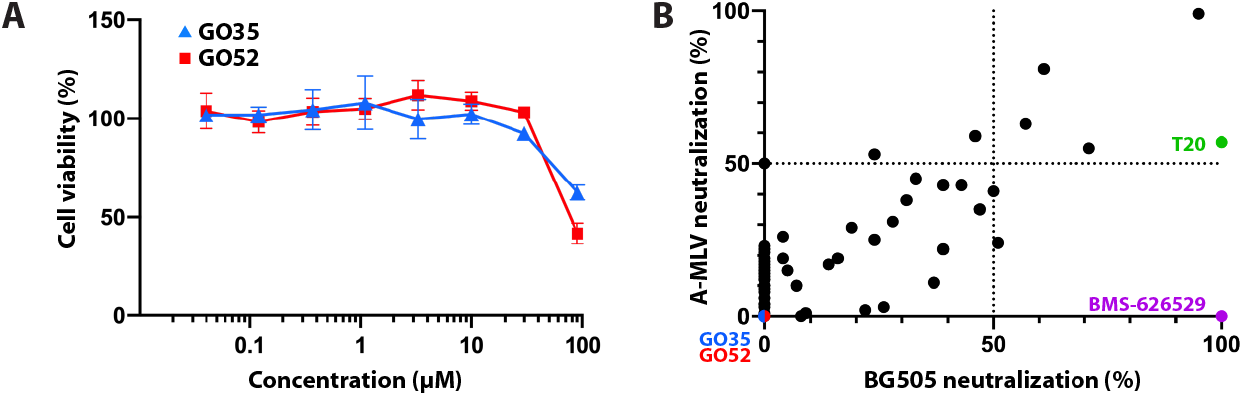
Cytotoxicity and neutralization assay profiles of small molecules. (**A**) Cytotoxicity measurements of GO35 and GO52. Assays performed as duplicates. Mean values plotted with standard deviation represented by vertical bars. (**B**) Neutralization activity of 80 small molecules against BG505 N332 HIV-1 and A-MLV viruses. All molecules tested at 30 μM final concentration. Assay performed as duplicates (*N=2*) and mean value is plotted. BMS-626529 and T20 (enfuvirtide) are known HIV-1 fusion inhibitors included as controls.

**Fig. S5.**
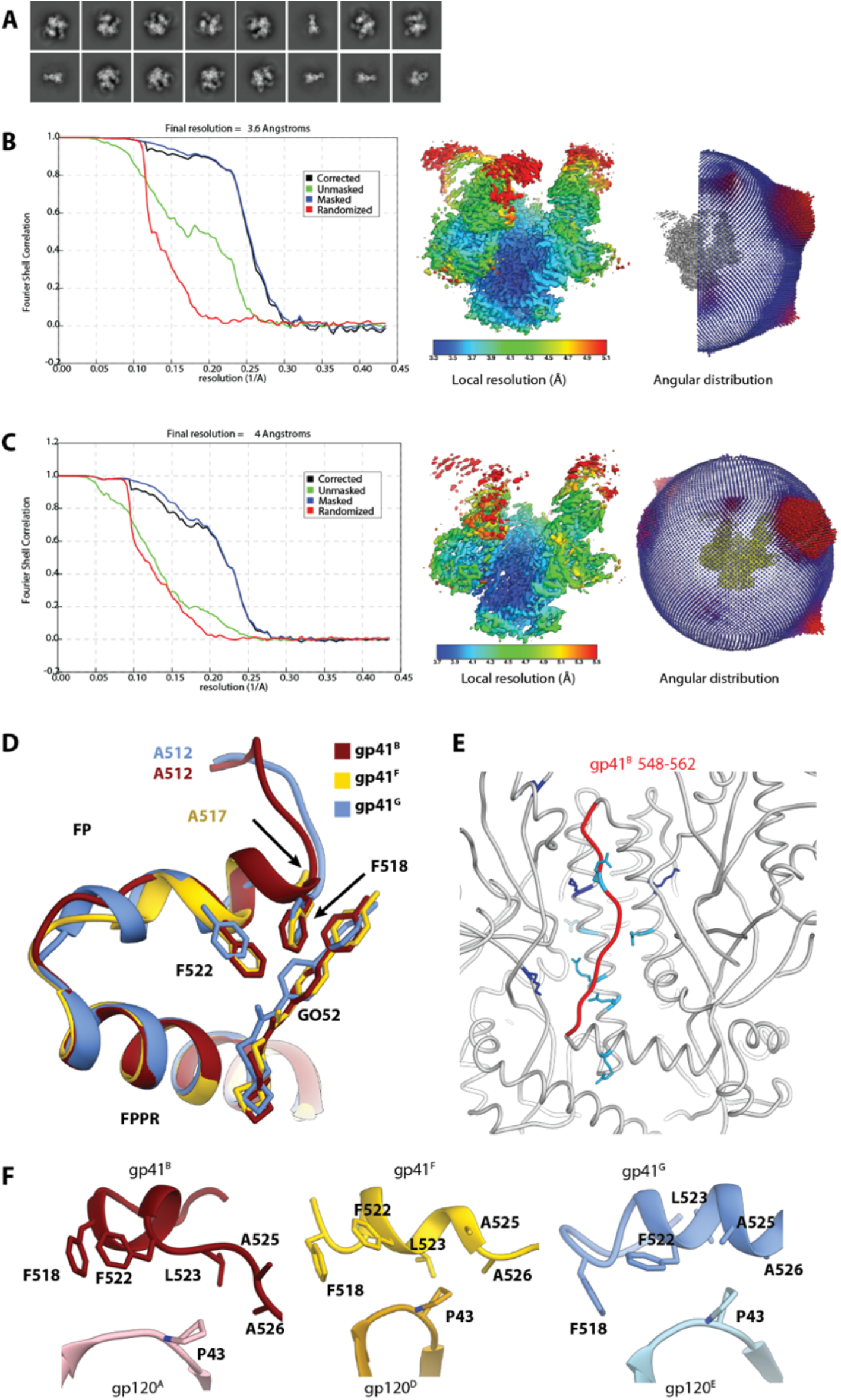
Cryo-EM reconstructions of B41+CD4+17b+GO52. (**A**) Select 2D class averages of B41+CD4+17b+GO52. Global Fourier shell correlations (FSC; *left*), local resolution estimates (colored by resolution in Å; *middle*) and distribution plot of particle orientations (*right*), of the (**B**) C3 symmetric and (**C**) asymmetric reconstructions of B41+CD4+17b+GO52. (**D**) Superimposition of the 3 asymmetric gp41 chains of (**C**) with a focus on the FP and FPPR regions. (**E**) Lysine (dark blue) and arginine (light blue) residues within 10 Å of gp41 HR1 region 548-562. (**F**) Comparison of P43 (gp120) interactions with the FP in the three asymmetric protomers of (**C**).

**Fig. S6.**
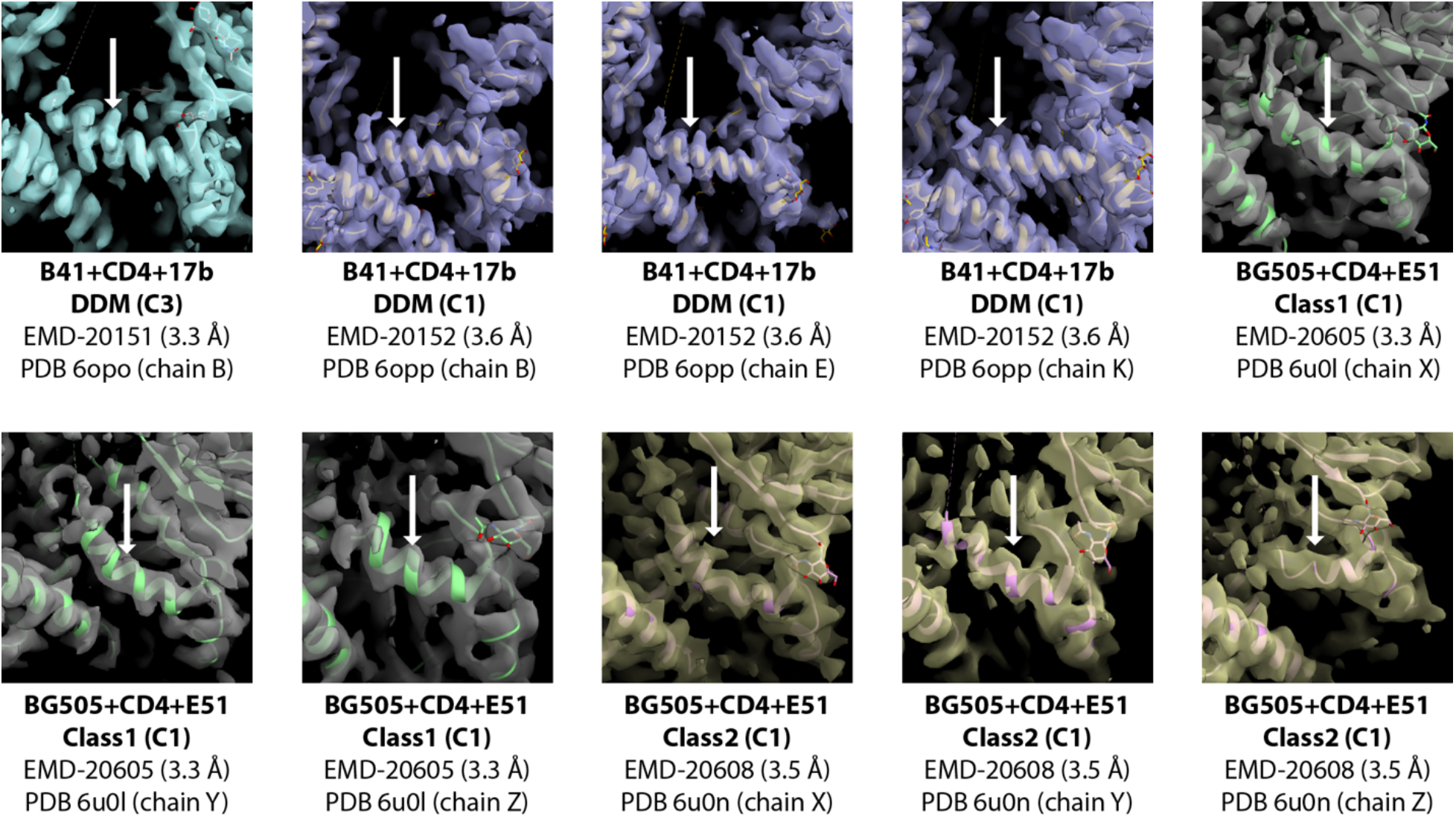
The B41 binding FPPR is more ordered than in BG505. Comparison of CD4- and DDM-bound C3-symmetric and asymmetric B41 cryo-EM maps with asymmetric reconstructions of CD4-bound BG505, with special focus on the FPPR (denoted by a white arrow).

**Fig. S7.**
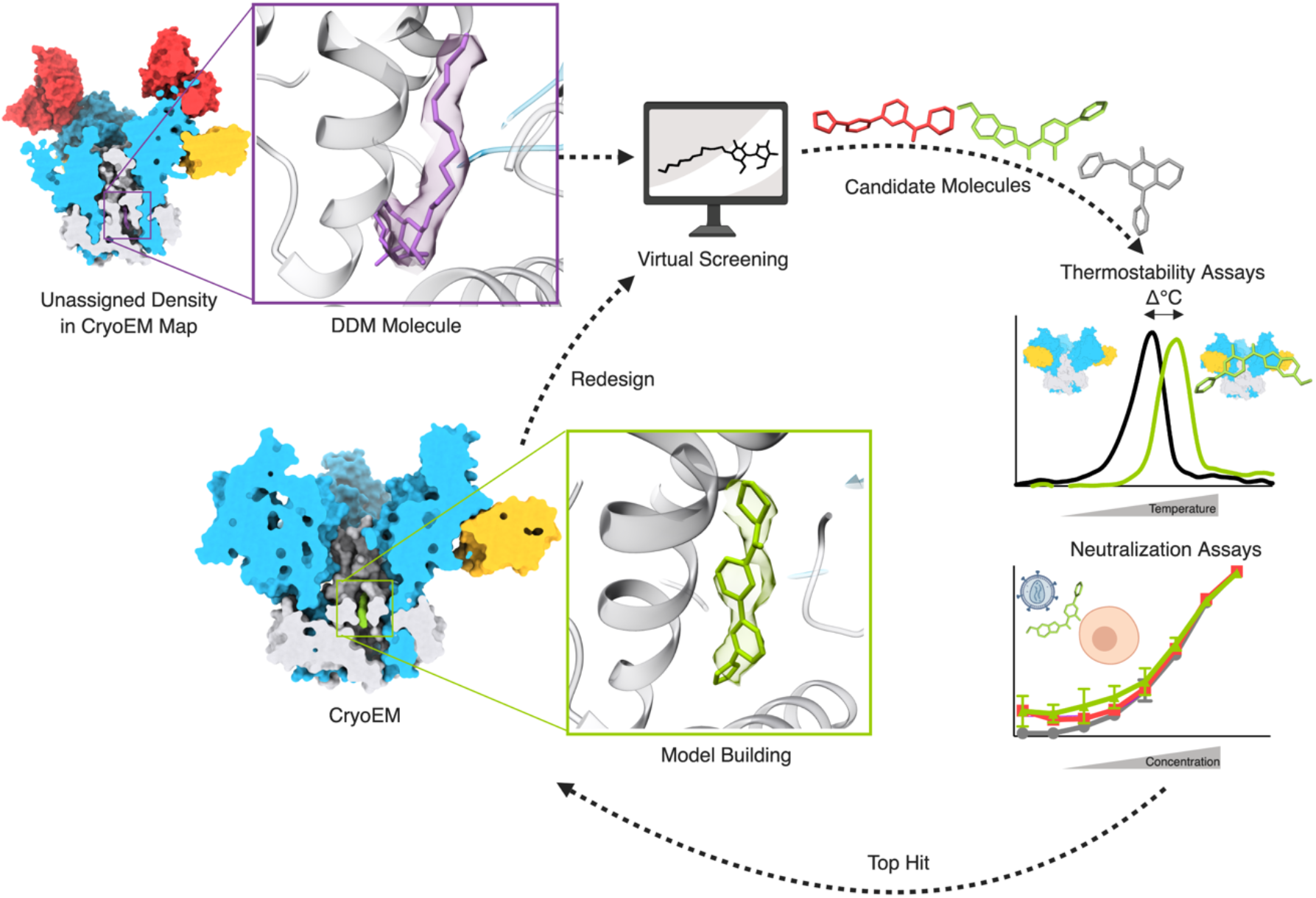
A schematic of iterative drug design using a combination of cryo-EM, virtual screening, and assays. Created with BioRender.com.

**Table S1.**
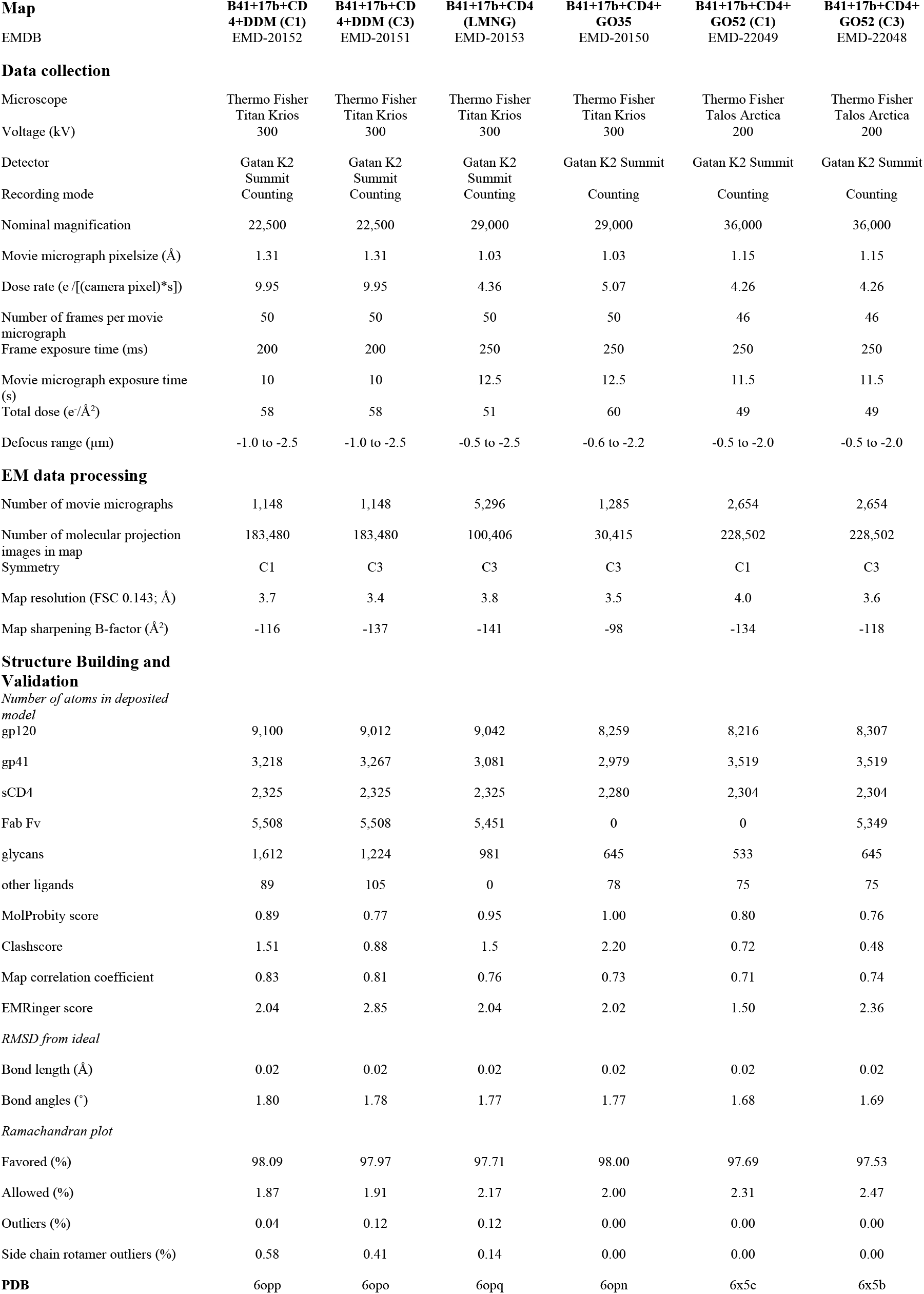
Cryo-EM data collection and modeling statistics.

**Table S2.** Difference in thermal stability of CD4-bound SOSIP trimers in the presence of small molecules relative to no small molecule controls as measured by differential scanning fluorimetry. Changes ≥ +1°C colored green, ≤ −1°C colored red. Absent values marked with a star denote that the small molecule had a high level of intrinsic fluorescence that interfered with the method.

| Small Molecule | ZINC ID | AMC011+CD4 | BG505+CD4 | B41+CD4 | CZA97+CD4 | DU422+CD4 | JRFL+CD4 | Small Molecule | ZINC ID | AMC011+CD4 | BG505+CD4 | B41+CD4 | CZA97+CD4 | DU422+CD4 | JRFL+CD4 |
| --- | --- | --- | --- | --- | --- | --- | --- | --- | --- | --- | --- | --- | --- | --- | --- |
| GO1 | ZINC00185349 | -0.1 | 0.1 | -0.4 | -0.1 | 0.6 | 0 | GO31 | ZINC67459760 | -0.4 | -0.4 | -0.3 | -0.1 | -0.3 | 0 |
| GO2 | ZINC01950659 | -0.1 | -0.1 | -0.3 | -0.5 | 0.6 | -0.2 | GO32 | ZINC77300595 | * | * | * | * | * | * |
| GO3 | ZINC00235832 | -0.1 | 0 | -0.3 | -0.3 | 0.7 | -0.3 | GO33 | ZINC67491033 | 0.1 | -0.1 | -0.1 | 0.1 | 0 | 0 |
| GO4 | ZINC00291882 | -0.1 | 0.2 | -0.3 | -0.3 | 1 | -0.2 | GO34 | ZINC97244092 | -0.5 | -0.4 | -1.1 | -1 | 1.2 | -0.8 |
| GO5 | ZINC00233220 | -0.1 | -0.1 | -0.3 | -0.3 | 1.1 | -0.1 | GO35 | ZINC96132694 | -0.6 | -1.1 | -3.1 | -1.5 | -1.1 | -1.9 |
| GO6 | ZINC01907589 | 0.1 | 0.1 | -0.3 | 0 | 1.1 | -0.2 | GO36 | ZINC91325610 | -0.4 | -0.4 | -0.7 | -0.8 | 0.3 | -0.8 |
| GO7 | ZINC03998067 | 0 | 0.1 | -0.3 | -0.2 | 0.6 | -0.2 | GO37 | ZINC67675494 | 0 | 0.2 | -0.4 | -0.6 | 0.8 | -0.2 |
| GO8 | ZINC00258817 | 0 | 0.1 | -0.5 | 0 | 0.4 | -0.2 | GO38 | ZINC20782685 | 0 | -0.5 | -0.7 | -0.6 | -0.4 | -0.5 |
| GO9 | ZINC00035601 | 0.2 | 0 | 0 | -0.1 | 0 | -0.2 | GO39 | ZINC65394371 | -0.2 | -0.1 | -0.2 | 0 | -0.2 | -0.1 |
| GO10 | ZINC12362428 | 0.1 | 0.1 | -0.2 | -0.1 | 0.4 | -0.1 | GO40 | ZINC65394381 | * | * | * | * | * | * |
| GO11 | ZINC00123118 | 0.2 | 0 | -0.4 | -0.1 | 0.5 | 0 | GO41 | ZINC55153733 | 0.2 | 0 | -0.9 | -0.6 | 0.8 | -0.4 |
| GO12 | ZINC18247429 | 0.1 | 0.1 | -0.2 | -0.2 | 0.5 | 0.2 | GO42 | ZINC71745481 | -0.6 | -0.7 | -1.1 | -0.4 | -0.6 | -0.7 |
| GO13 | ZINC00074920 | 0.1 | -0.1 | -0.2 | -0.1 | 0.5 | 0.1 | GO43 | ZINC95366696 | 0.1 | 0.2 | -0.4 | -0.4 | 1.4 | -0.2 |
| GO14 | ZINC00569594 | -0.1 | 0.1 | -0.2 | -0.4 | 1.3 | -0.2 | GO44 | ZINC77386563 | * | * | * | * | * | * |
| GO15 | ZINC00461274 | -0.3 | -0.1 | -0.2 | -0.1 | -0.3 | -0.1 | GO45 | ZINC67803542 | 0 | 0.1 | -0.3 | -0.4 | 1.1 | -0.2 |
| GO16 | ZINC00462885 | 0.1 | 0.2 | -0.2 | -0.1 | 0.4 | 0 | GO46 | ZINC77389895 | -0.3 | -0.4 | -1 | -0.6 | 0.3 | -0.7 |
| GO17 | ZINC00463378 | * | * | * | * | * | * | GO47 | ZINC96149848 | * | * | * | * | * | * |
| GO18 | ZINC00469391 | -0.2 | -0.1 | 0.1 | 0.1 | 0.2 | -0.1 | GO48 | ZINC55252643 | -0.2 | 0.1 | -0.5 | -0.5 | 0.1 | -0.2 |
| GO19 | ZINC04263843 | 0.8 | -0.2 | -0.2 | 0.1 | 1 | -0.2 | GO49 | ZINC65422611 | -0.1 | 0 | -0.1 | -0.2 | -0.3 | -0.1 |
| GO20 | ZINC05059895 | -1.3 | -0.9 | -0.7 | -1.2 | -1.3 | -0.8 | GO50 | ZINC14974795 | -0.1 | -0.1 | -0.7 | -0.6 | 0.1 | -0.2 |
| GO21 | ZINC00621192 | 0.2 | 0 | -0.1 | 0 | 0.8 | -0.4 | GO51 | ZINC65455118 | 0.1 | 0.4 | -0.6 | 0 | 0.2 | -0.1 |
| GO22 | ZINC05061043 | 0 | -0.2 | -0.5 | -0.2 | 1.1 | 0 | GO52 | ZINC97681845 | -0.3 | 0 | -0.4 | -0.7 | 1.3 | -0.2 |
| GO23 | ZINC04247959 | 0.1 | 0.2 | -0.1 | -4.3 | 2.4 | 0 | GO53 | ZINC91819525 | -0.1 | -0.2 | -0.8 | -0.6 | 0.8 | -0.5 |
| GO24 | ZINC00541458 | 0.2 | 0.2 | -0.3 | -0.1 | 1.1 | 0.2 | GO54 | ZINC67843155 | -0.4 | -0.3 | -0.6 | -0.2 | -0.2 | -0.3 |
| GO25 | ZINC01428810 | 0.2 | 0.2 | -0.5 | -0.1 | 0.8 | 0.1 | GO55 | ZINC97766602 | -0.2 | -0.2 | -0.1 | -0.5 | 0 | -0.3 |
| GO26 | ZINC10461636 | 0.2 | 0.3 | -0.1 | -0.4 | 2 | 0.2 | GO56 | ZINC72171577 | -0.4 | -0.3 | -0.5 | -0.4 | -0.2 | -1.8 |
| GO27 | ZINC06767131 | 0.3 | 0.2 | -0.2 | -0.2 | 2.1 | -0.1 | GO57 | ZINC67975161 | -0.7 | -1 | -1 | -1.2 | -2 | -1.6 |
| GO28 | ZINC36351253 | * | * | * | * | * | * | GO58 | ZINC67975538 | * | * | * | * | * | * |
| GO29 | ZINC36358921 | 0.2 | 0.1 | -0.1 | -0.1 | 1.9 | 0 | GO59 | ZINC72173974 | -0.1 | -0.4 | -0.6 | -0.7 | -0.2 | -0.4 |
| GO30 | ZINC77286437 | * | * | * | * | * | * |  |  |  |  |  |  |  |  |

